# FT-GO: a multiplex fluorescent tyramide signal amplification system for histochemical analysis

**DOI:** 10.1101/2022.07.22.501050

**Authors:** Kenta Yamauchi, Shinichiro Okamoto, Yoko Ishida, Kohtarou Konno, Kisara Hoshino, Takahiro Furuta, Megumu Takahashi, Masato Koike, Kaoru Isa, Masahiko Watanabe, Tadashi Isa, Hiroyuki Hioki

## Abstract

Tyramide signal amplification (TSA) is a highly sensitive method for histochemical analysis. Previously, we reported a TSA system, BT-GO, for bright-filed imaging. Here, we develop FT-GO (Fluorochromized Tyramide-Glucose Oxidase) as a multiplex fluorescent TSA system. FT-GO involves peroxidase-catalyzed deposition of FT with hydrogen peroxide produced in enzymatic reaction between glucose and glucose oxidase. We showed that FT-GO enhanced immunofluorescence signals while maintaining low background signals. Compared with indirect immunofluorescence detections, FT-GO demonstrated a more widespread distribution of monoaminergic projection systems in mouse and marmoset brains. For multiplex labeling with FT-GO, we quenched Ab-conjugated peroxidase using sodium azide. We applied FT-GO to multiplex fluorescent *in situ* hybridization, and succeeded in labeling neocortical interneuron subtypes by coupling with immunofluorescence. FT-GO immunofluorescence further increased the detectability of an adeno-associated virus tracer. Given its simplicity and a staining with a high signal-to-noise ratio, FT-GO would provide a versatile platform for histochemical analysis.

## Introduction

Tyramide signal amplification (TSA), also called catalyzed reporter deposition (CARD), is a highly sensitive enzymatic method that enables the detection of low-abundance targets in histochemical analysis. TSA utilizes the catalytic activity of peroxidase (POD) to yield high-density labeling of targets *in situ*^1,2^. POD reacting with hydrogen peroxide (H_2_O_2_) catalyzes oxidative condensation of tyramide to produce highly reactive intermediates, tyramide radicals. The activated tyramide covalently binds to electron-rich moieties such as tyrosine, phenylalanine and tryptophan at or near the POD. The short half-life of the tyramide radicals restricts the deposition of tyramide^3^.

TSA was originally introduced for the detection of immunosorbent and immunoblotting assays more than thirty years ago^1^. Soon after its introduction, TSA was adapted to immunohistochemistry (IHC), lectin histochemistry and neuroanatomical tract tracing methods by Adams^4^. TSA has been further implemented in detection procedures for DNA and RNA *in situ* hybridization (ISH) histochemistry^5-7^ and electron microscopy^8,9^. Given its simplicity, high sensitivity, high specificity and broad applicability, TSA has become an essential tool for histochemical analysis.

Previously, we reported a straightforward and cost-effective TSA system, BT-GO (Biotinyl Tyramine-Glucose Oxidase), for bright-filed imaging^10,11^. Unlike conventional TSA systems, where H_2_O_2_ is added directly to their reaction mixtures^4-7,12^, BT-GO utilizes H_2_O_2_ produced by oxidation of glucose by glucose oxidase to deposit BT onto tissues^13^. H_2_O_2_ is thermodynamically unstable and readily decomposes to oxygen and water in light, heat or in the presence of metal ions. Enzymatic reaction between glucose and glucose oxidase supplies the compound stably during BT-GO reaction. BT-GO provides a substantial increase in sensitivity; we^10,11^ and others^14,15^ have applied BT-GO to chromogenic IHC including axonal projection analysis. BT-GO has been further adapted for an mRNA fluorescence ISH (FISH) technique by coupling with fluorochrome-conjugated streptavidin^16-18^. Although haptenized tyramides including BT are invaluable reagents for TSA, fluorochromized tyramide (FT) can be visualized directly by fluorescent microscopy without additional staining with avidin or antibodies (Abs) against haptens, which potentially induce background signals. FT further permit multiplex histochemical labeling that allows for simultaneous detection of multiple targets within a single specimen, enabling analysis of spatial organization of the target molecules in the context of the tissue architecture.

Here, we developed FT-GO (Fluorochromized Tyramide-Glucose Oxidase) as a multiplex fluorescent TSA system. FT-GO involves POD-catalyzed deposition of FT with H_2_O_2_ produced during oxidation of glucose by glucose oxidase, as with BT-GO. We adapted FT-GO to immunofluorescence (IF), multiplex IF, multiplex mRNA FISH histochemistry and neuroanatomical tract tracing with an adeno-associated virus (AAV) tracer. FT-GO provided strong labeling with a high signal to noise ratio in each application.

## Results

### Signal enhancement in IF with FT-GO

FT-GO is a fluorescent TSA system that utilizes H_2_O_2_ produced during oxidation of glucose by glucose oxidase to deposit FT onto tissues (Fig. 1). H_2_O_2_ is thermodynamically unstable and decomposes to form water and oxygen. In the FT-GO signal amplification, enzymatic reaction between glucose and glucose oxidase exerts stable supply of the thermodynamically unstable compound. The phenolic part of FT and H_2_O_2_ react with POD to produce highly reactive intermediates, tyramide radicals, which in turn bind to electron-rich moieties such as tyrosine residue in a covalent manner in the vicinity of POD.

**Fig. 1.**
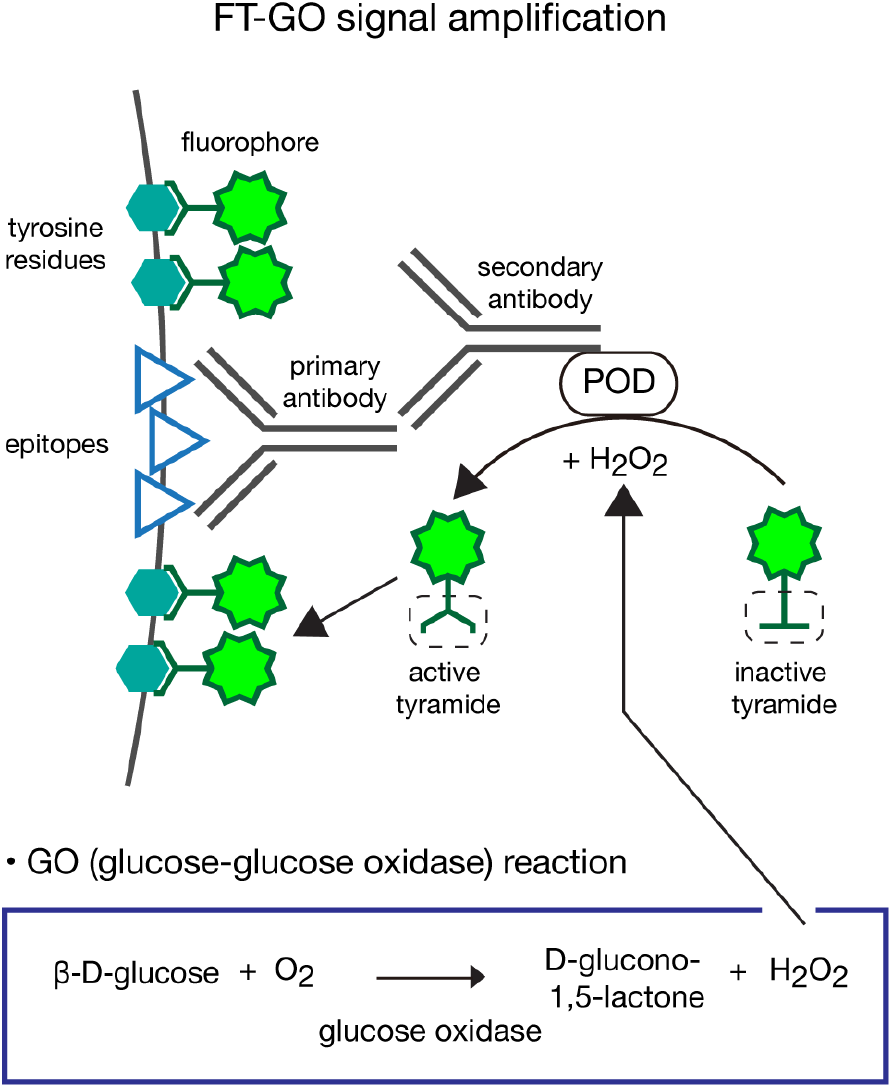
FT-GO signal amplification. Schematic diagram of FT-GO signal amplification in IF. FT-GO is a fluorescent TSA system that utilizes H_2_O_2_ produced during oxidation of glucose by glucose oxidase to deposit FT. POD reacts with the H_2_O_2_, and oxidizes the phenolic part of tyramide to produce highly reactive intermediates, tyramide radicals, which in turn covalently bind to electron-rich moieties such as tyrosine residues at or near the POD.

We first examined signal amplification characteristics of FT-GO with IF for a neuronal marker, Rbfox3 (also known as neuronal nuclei [NeuN]). Using direct, indirect and FT-GO IF methods, we stained layer 2/3 neurons in the mouse primary somatosensory cortex (S1) with an Ab against Rbfox3 (clone A60^19^). Brain sections containing the S1 were reacted with the Ab serially diluted 10-fold from 1:100 to 1:100, 000 (Fig. 2). Fluorescence signals amplified with FT-GO were much stronger than those detected with the direct and indirect IF detections at any concentrations tested (Fig. 2a-l). Quantitative analysis showed that FT-GO yielded 64.4 to 183.4 fold and 9.7 to 34.1 fold signal amplification compared with the direct and indirect IF detection, respectively (Fig. 2m). The fluorescence signals obtained by FT-GO were less affected by Ab dilution: the ratio of maximum to minimum mean signal intensity in the serial dilution experiments was 3.1, 3.9 and 1.1 for the direct, indirect and FT-GO IF detections.

**Fig. 2.**
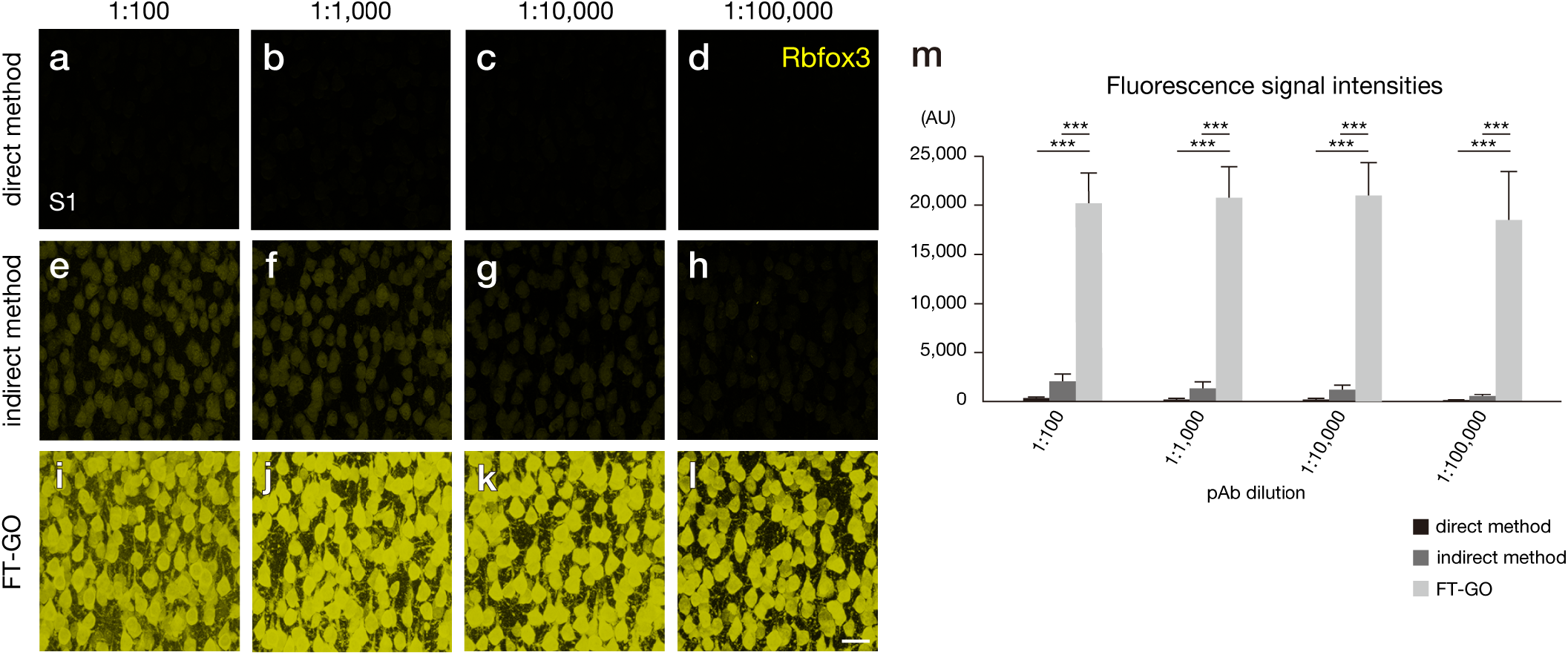
Signal amplification in IF with FT-GO. **a-l**, IF for Rbfox3 using a direct (**a-d**), indirect **(e-h**) and FT-GO method (**i-l**) in layer 2/3 neurons of the mouse S1 (n = 3 animals for each condition). Brain sections were reacted with an anti-Rbfox3 Ab at dilutions of 1:100 (**a, e, i**), 1:1,000 (**b, f, j**), 1;10,000 (**c, g, k**) and 1:100,000 (**d, h, l**). CF®488A tyramide is used for color development in the FT-GO method. Images are acquired with the same parameters for comparisons. **m**, Histograms representing fluorescence signal intensities (AU) of Rbfox3-positive cells in the layer 2/3 of S1 (n = 630 cells, pAb 1:100, direct method; n = 644 cells, pAb 1:100, indirect method; n = 686 cells, pAb 1:100, FT-GO; n = 622 cells, pAb 1:1,000, direct method; n = 698 cells, pAb 1:1,000, indirect method; n = 758 cells, pAb 1:1,000, FT-GO; n = 620 cells, pAb 1:10,000, direct method; n = 639 cells, pAb 1:10,000, indirect method; n = 728 cells, pAb 1:10,000, FT-GO; n = 650 cells, pAb 1:100,000, direct method; n = 619 cells, pAb 1:100,000, indirect method; n = 686 cells, pAb 1:100,000, FT-GO; n = 3 animals for each condition, pAb 1:100; *H* □ = 1737, *df*□ =C2, *P* < 0.0001, Kruskal–Wallis test, pAb 1:1,000, *H* □ = 1832, *df*□ = □2, *P* < 0.0001, Kruskal–Wallis test, pAb 1:10,000; *H* □ = 1744, *df*□ = □2, *P* < 0.0001, Kruskal–Wallis test, pAb 1:100,000; *H* □ = 1726, *df*□ = □2, *P* < 0.0001, Kruskal–Wallis test, *** *P* < 0.0001; Dunn’s *post hoc* test). Data are represented as means ± SDs. Scale bar: 25 µm.

Monoaminergic fibers, such as catecholaminergic (CA) and serotonergic (5-HT) fibers, are highly ramified into fine varicose axonal processes in their projection areas^20,21^, hampering high-fidelity visualization of the projection systems. We used FT-GO signal amplification to visualize the fine axonal morphology of monoaminergic fibers. Tyrosine hydroxylase (TH) and serotonin transporter (Slc6a4) were chosen as makers for CA and 5-HT fibers. Compared with an indirect IF detection, FT-GO showed a more diffuse and widespread distribution of TH-immunoreactivity (IR) throughout the brain, in addition to the dense and localized IR in brain regions such as the caudate-putamen (CPu), amygdala and hypothalamus (Fig. 3a, e). At high magnification, FT-GO demonstrated fine axonal morphology of TH-positive fibers in the cerebral cortex, hippocampus and thalamus, where TH-positive fibers diffusely innervated (Fig. 3f-h). In contrast, TH-positive fibers were only faintly labeled in these brain regions using the indirect detection (Fig. 3b-d). FT-GO also provided dramatically increased sensitivity in Slc6a4 IF: while FT-GO visualized a diffuse and widespread projection of Slc6a4-positive fibers throughout the brain, only faintly labeled fibers were observed in indirect IFs under the same imaging condition (Supplementary Fig. S1a, b). FT-GO further succeeded in visualizing 5-HT axons at single fiber resolution in brain regions including the cerebral cortex, hippocampus and thalamus (Supplementary Fig. S1c-e). Importantly, in the IF assay, we used five times lower concentration of the primary Ab in the FT-GO detection in comparison with the indirect detection.

**Fig. 3.**
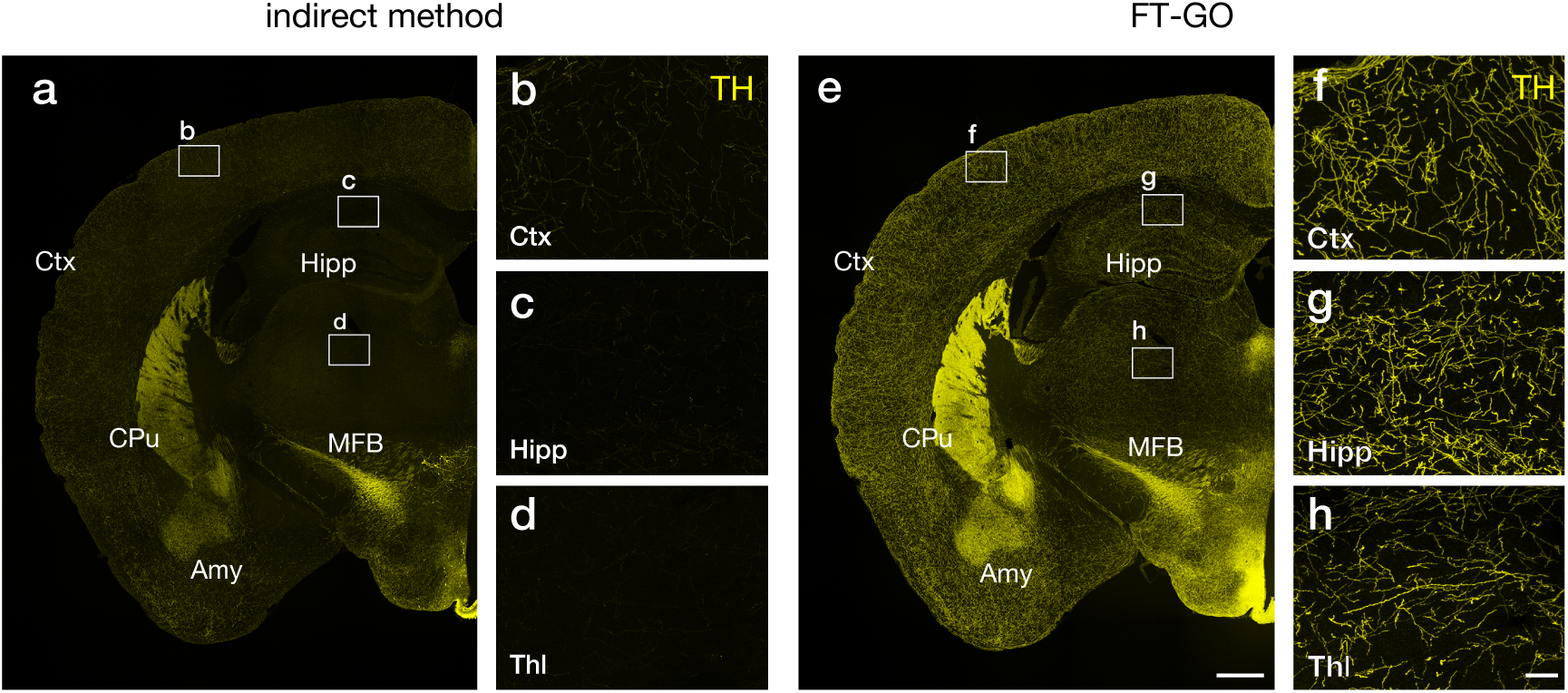
CA fiber innervation in the mouse brain visualized with FT-GO. **a-h**, TH IF in mouse brain sections visualized with an indirect (**a-d**) and FT-GO method (**e-h**) (n = 3 animals for each condition). (**b-d)** and (**f-h)** show higher magnification images in rectangles in (**a**) and (**e**). CF®488A tyramide is used for color development in the FT-GO method. Amy: amygdala, CPu: caudate-putamen, Ctx: cerebral cortex, Hipp: hippocampus, MFB: medial forebrain bundle, Thl: thalamus. Scale bars: 500 µm in (**e**) and 50 µm in (**h**).

We then explored the applicability and scalability of FT-GO signal amplification to large-brained animals. The common marmoset (*Callithrix jacchus*) is becoming increasingly popular as a non-human primate model for neuroscience research^22,23^. We adapted FT-GO to IF microscopy experiments with an Ab against TH in the marmoset brain. FT-GO again provided an increase in sensitivity in brain sections of marmosets, compared with an indirect IF detection (Fig. 4). While dense and localized IF signals for TH were detected in brain regions such as the caudate nucleus, putamen and hypothalamus in FT-GO IFs, only weak signals were obtained by the indirect IF under the same imaging condition (Fig. 4a, b). Additionally, FT-GO showed diffused IRs for TH in brain regions such as the cerebral cortex, thalamus and hippocampus (Fig. 4b). High power views revealed individual TH-positive fibers in these brain regions (Fig. 4c-e). We further found cell bodies frequently labeled for TH by FT-GO IF in the cerebral cortex (Fig. 4c), which have not been described in the marmoset.

**Fig. 4.**
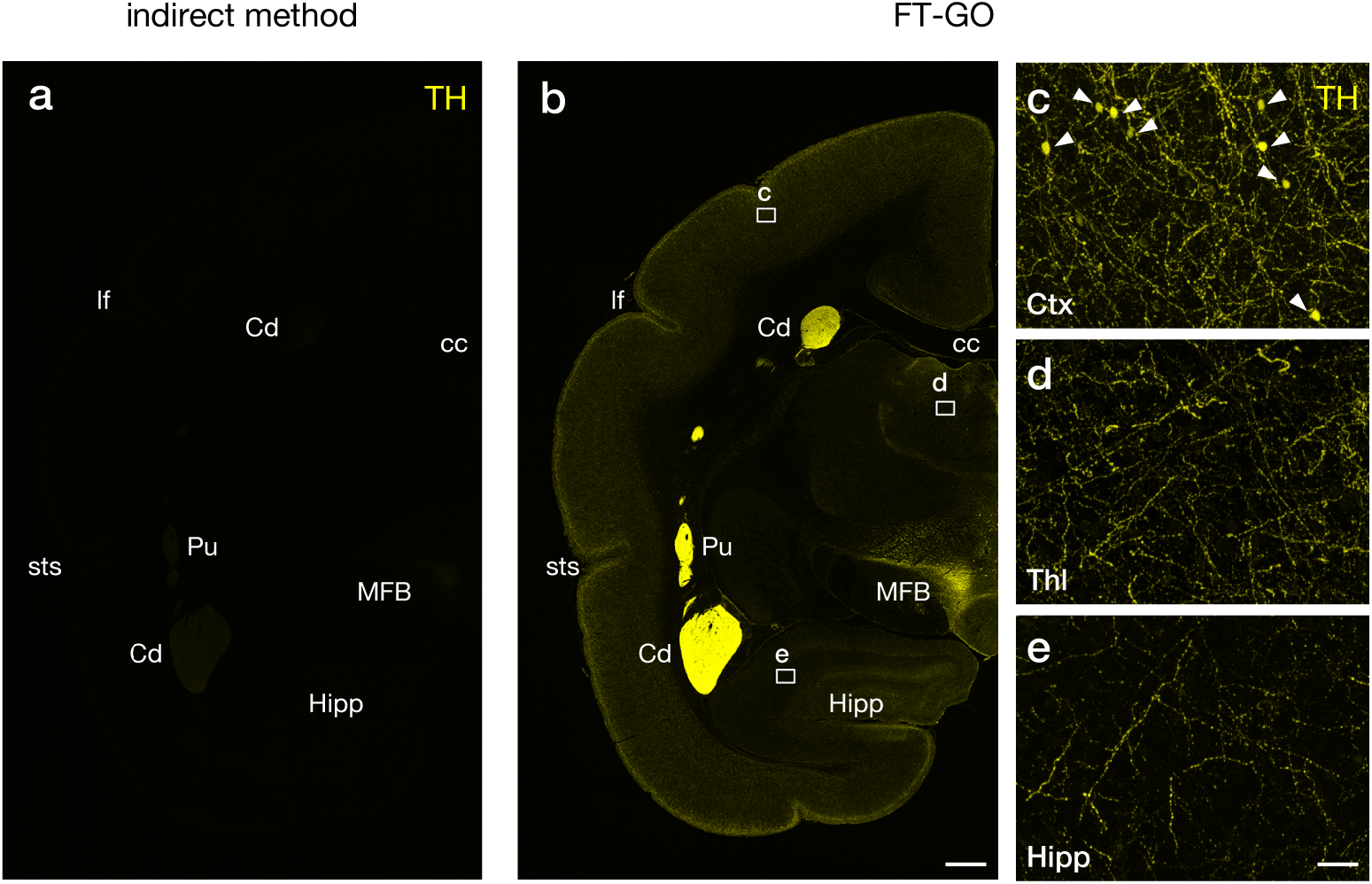
CA fiber innervation in the marmoset brain visualized with FT-GO. **a, b**, TH IF in marmoset brain sections visualized with an indirect (**a**) and FT-GO method (**b**) (n = 2 animals for each condition). Images are acquired with the same parameters for comparisons. **c-e**, Higher magnification images in rectangles in (**b**). Arrowheads in (**c**) indicate TH-positive cell bodies in the neocortex. CF®647 tyramide is used for color development in the FT-GO method. cc: corpus callosum, Cd: caudte nucleus, Ctx: cerebral cortex, Hipp: hippocampus, lf: lateral fissure, MFB: medial forebrain bundle, Pu: putamen, sts: superior temporal sulcus, Thl: thalamus. Scale bars: 1 mm in (**b**) and 50 µm in (**e**).

### Multiplex IF and mRNA FISH histochemistry with FT-GO

Simultaneous detection of multiple distinct targets within a single tissue section allows for analyzing spatial organization of the targets in the context of the tissue architecture. For adaptation of multiplex labeling with FT-GO, Ab-conjugated POD must be inactivated prior to subsequent detection rounds. A number of methods have been described for quenching POD activity, including incubation with H_2_O_2_, low pH buffer, and NaN_3_. Heat-mediated removal of primary/secondary Ab-POD complex has been implemented to TSA-based multiplex IF^24,25^. Of these, we choose NaN_3_ for quenching POD activity because of its less deleterious effects on antigenicity, antigen-antibody binding, hybridized strands and tissue integrity. We first asked whether NaN_3_ was effective at quenching Ab-conjugated POD in FT-GO. For this, we carried out two rounds of FT-GO with different color FT sequentially in IF for Rbfox3. Between the first and second round, brain sections were treated with high concentration of NaN_3_ (2% [w/v]) or phosphate buffered saline (PBS). Ineffective quenching of Ab-conjugated POD should result in deposit of the FT used in the second round of FT-GO due to residual activity of POD. We found effective inactivation of Ab-conjugated POD with NaN_3_: while fluorescence signals from the second round of FT-GO were detected in almost all cells labeled by the first round of FT-GO in PBS-treated brain sections (Supplementary Fig. S2a-c), these signals were completely vanished by incubation with NaN_3_ (Supplementary Fig. S2d-f). We then conducted multiplex IF with FT-GO. Mouse hippocampal sections were reacted with Abs against Rbfox3, an astrocytic marker, glial fibrillary acidic protein (Gfap) and a microglial marker, allograft inflammatory factor 1 (Aif1) (also known as ionized calcium binding adaptor molecule 1 [Iba1]), and these Abs were multiply detected with FT-GO. Specific distribution and no obvious colocalization of Rbfox3, Gfap and Aif1 IRs in the mouse hippocampal region demonstrated multiplex detection with FT-GO (Fig. 5a-i). FT-GO IF for Rbfox3 clearly showed pyramidal cell layers of cornu ammonis regions and granule cell layers of the dentate gyrus (DG) (Fig. 5a, d, h). FT-GO IFs for Gfap and Aif1 revealed near even distributions of astrocytic and microglial cells in the mouse hippocampal formation (Fig. 5a). At high magnification, we observed thick Gfap cytoskeleton radiating out from a central hub (Fig. 5b, f) and thin tortuous and ramified processes emanating from small cell bodies of microglial cells, which is characteristic of resting microglia (Fig. 5c, g). FT-GO signal amplification substantially increased Gfap IR, morphologically resembling reactive astrocytes (Fig. 5b, f). FT-GO IF further demonstrated tubular IRs for Gfap, which was likely astrocytic endfeet surrounding blood vessels (Fig. 5b), and Gfap-positive processes oriented radially in the supra- and infra-blades of DG (Fig. 5f). The Gfap-positive processes were extended from the subgranular zone of DG that was determined by FT-GO IF for Rbfox3 (Fig. 5f, i), indicating Gfap-positive processes emanating from neural stem cells^26^.

**Fig. 5.**
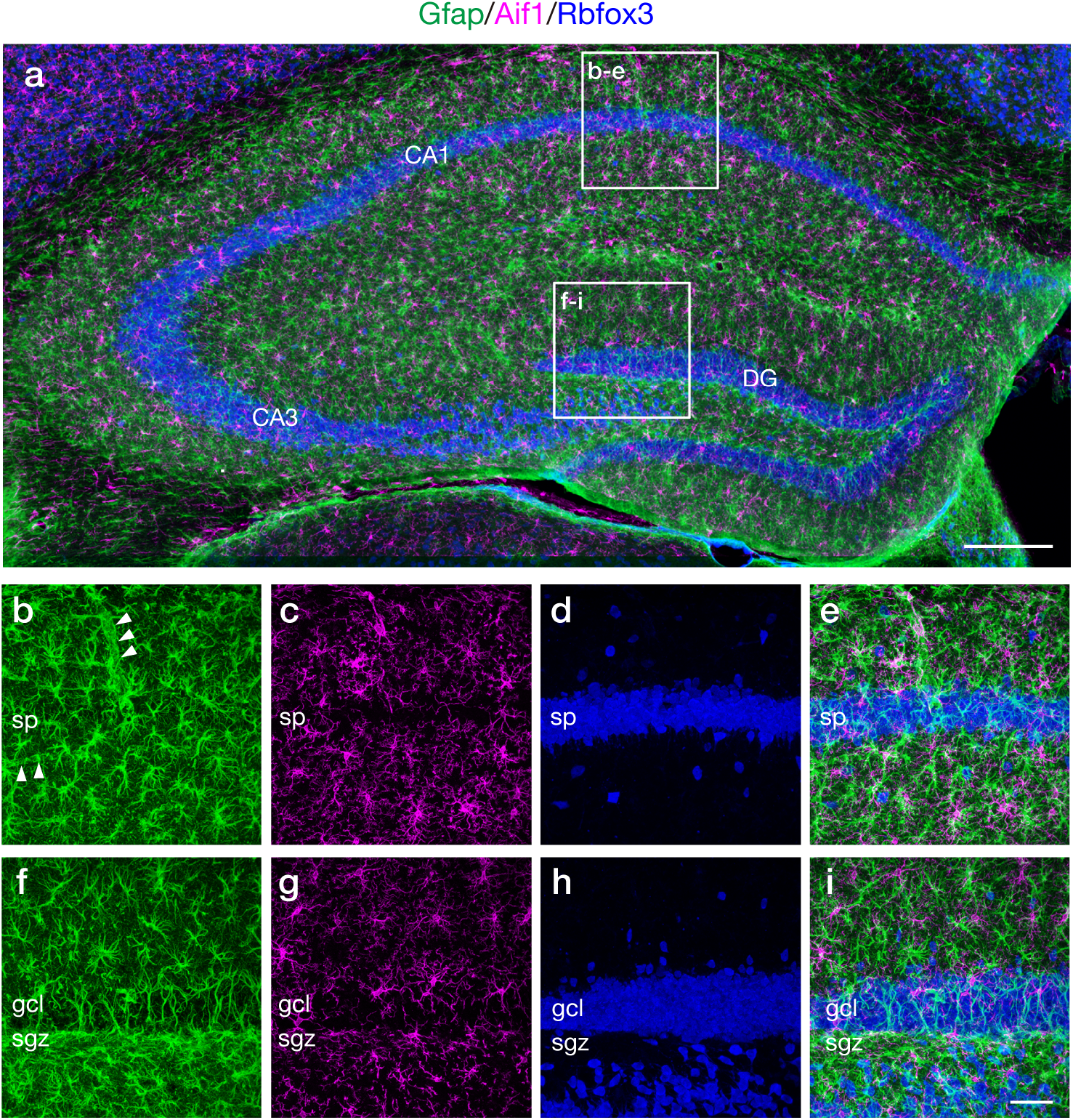
Fluorescence triple labeling with multiplex FT-GO IF. **a**, Gfap (green), Aif1 (magenta) and Rbfox3 (blue) multiplex FT-GO IF in the mouse hippocampus (n = 3 animals). **b-i**, Higher magnification images in rectangles in (**a**). **b-d, f-h**, Representative images of triple labeling for Gfap (**b, f**), Aif1 (**c, g**) and Rbfox3 (**d, h**). **e, i**, Triple merged images of (**b-d**) and (**f-h**). CF®488A, CF®568 and CF®647 tyramide are used for color development of Gfap, Aif1 and Rbfox3 IRs, respectively. Arrowheads indicate tubular IR for Gfap. Scale bars: 200 µm in (**a**) and 50 µm in (**i**).

Multiplex mRNA FISH histochemistry facilitates identifying molecular signatures of individual cell populations in fixed tissues^27-29^. Neocortical interneurons contain a tremendous diversity in term of their gene expressions^30-32^. We then adapted FT-GO to multiplex mRNA FISH histochemistry, and combined with indirect IF detections for neurochemical markers to label specific subtypes of interneurons. We focused on parvalbumin (Pvalb) or cholecystokinin (Cck)-expressing GABAergic interneurons that compose two major populations of perisomatic interneurons in the mouse cerebral cortex^33^. We performed fluorescence quadruple labeling experiments for Slc17a7 (also known as vesicular glutamate transporter 1 [Vglut1]) mRNA, Gad1 (also known as glutamate decarboxylase 67 [Gad67]) mRNA, Pvalb protein and proCck peptide to label Pvalb or Cck-expressing neocortical GABAergic interneurons. *Slc17a7* and *Gad1* are markers for glutamatergic and GABAergic neurons in the dorsal telencephalon. Cck is also expressed by subpopulations of neocortical glutamatergic neurons^34^. Mouse brain sections were hybridized with complementary RNA (cRNA) probes for *Slc17a7* and *Gad1*, and FT-GO signal amplification was applied in the detection procedure. The Ab-POD conjugated was inactivated with NaN_3_ after each round of FT deposition. Hybridization signals for *Slc17a7* and *Gad1* were clearly visualized with FT-GO (Fig. 6a-c): while *Slc17a7* cRNA probe densely labeled all layers of the cerebral cortex, pyramidal cell layers of CA regions and granule cell layers of the DG, scattered hybridization signals of *Gad1* were found in these brain regions. IF signals for Pvalb and Cck were also clearly detected in our labeling system (Fig. 6d, e). We succeeded in labeling Pvalb and Cck-expressing neocortical GABAergic interneurons: neurons that expressed Gad1 mRNA and Pvalb protein, and Gad1 mRNA and proCck peptide were found in the quadruple labeling (Fig. 6g-i). Mutually exclusive expression of *Slc17a7* and *Gad1* validated multiplex detection of mRNA expressions with FT-GO (Fig. 6f).

**Fig. 6.**
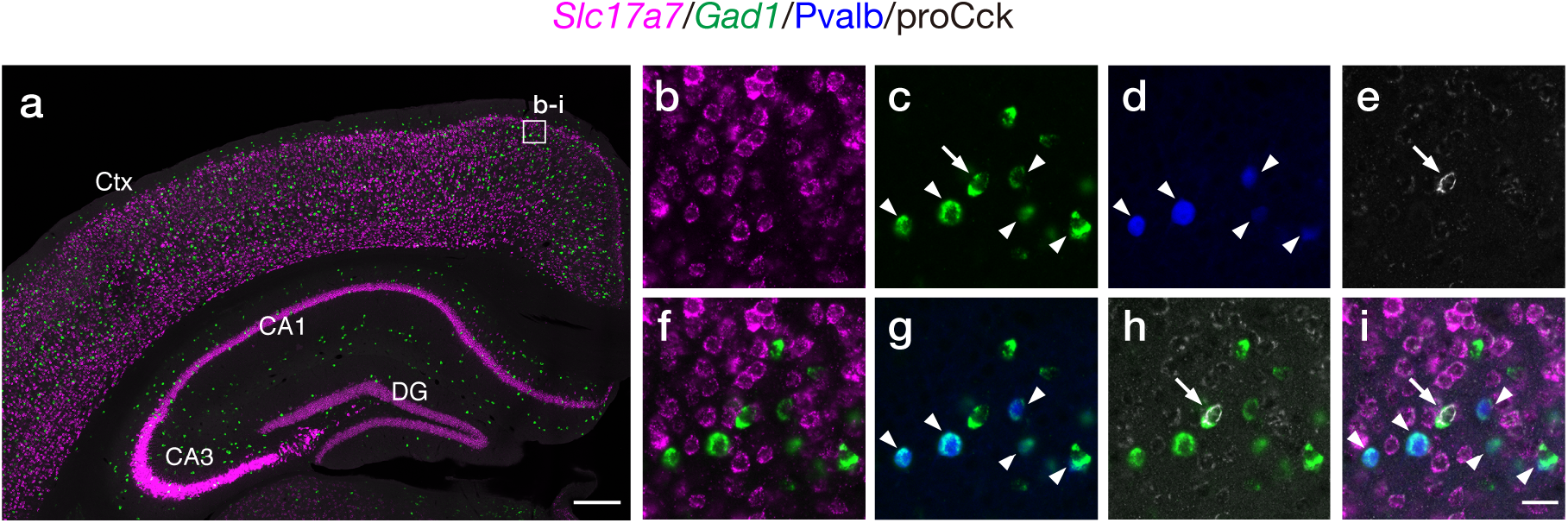
Fluorescence quadruple labeling with multiplex FT-GO FISH and IF. **a**, *Slc17a7* (magenta) and *Gad1* (green) FT-GO mRNA FISH in the mouse brain (n = 3 animals). **b-e**, Representative images of quadruple labeling for Slc17a7 mRNA (**b**), Gad1 mRNA (**c**), Pvalb protein (blue, **d**) and proCck peptide (gray, **e**) in the rectangle in (**a**). **f-h**, Double merged images of *Slc17a7* and *Gad1* (**f**), *Gad1* and Pvalb (**g**), and *Gad1* and proCck (**h**). **i**, A quadruple merged image of *Slc17a7, Gad1*, Pvalb, and proCck. CF®488A and CF®568 tyramide are used for color development of *Gad1* and *Slc17a7* hybridization signals. Pvalb- and Cck-positive GABAergic interneurons are indicated by arrowheads and arrows, respectively. Scale bars: 300 µm in (**a**) and 25 µm in (**i**).

### Increase of detectability of an AAV tracer with FT-GO IF

AAV vectors carrying fluorescent protein (FP) reporter genes are widely used as anterograde neuronal tracers with equivalent sensitivity and specificity to classical tracers^35,36^. Although AAV vectors mediate robust fluorescence labeling of infected neurons, Ab amplification for the FPs is useful to increase detectability of AAV tracers^37,38^. TSA has been adapted to neuroanatomical tract tracing studies with classical tracers^4^, and FT-GO increased detectability of IF compared with indirect detections (Fig. 2-4, Supplementary Fig. S1). Lastly, we adapted FT-GO IF to increase detectability of an anterograde AAV tracer, AAV2/1-SynTetOff-mRFP1. AAV-SynTetOff is a single AAV vector Tet-Off platform which is composed of regulator and response elements separated by an insulator^39^. The regulator element expresses tTA, which is driven a human synapsin I promoter, whereas the response element produced a reporter protein under the control of a TRE promoter. The AAV tracer was stereotaxically injected into the CPu to label the striatofugal projection system. The AAV-injected mice were sacrificed seven to ten days after injections. Brain sections were cut along the striatonigral axis and adjacent sections were examined for fluorescence imaging of mRFP1 and FT-GO IF with an Ab against RFP (Fig. 7). FT-GO IF substantially increased detectability of the anterograde AAV tracer. While native fluorescence of mRFP1 was hardly detected outside the CPu (Fig. 7a), FT-GO IF visualized the whole striatofugal projection system: FT-GO IF labeled fibers arising from the CPu extended caudally to the brainstem, forming axon terminal fields in the external segment of the globus pallidus (GPe) and substantia nigra (SN) (Fig. 7b).

**Fig. 7.**
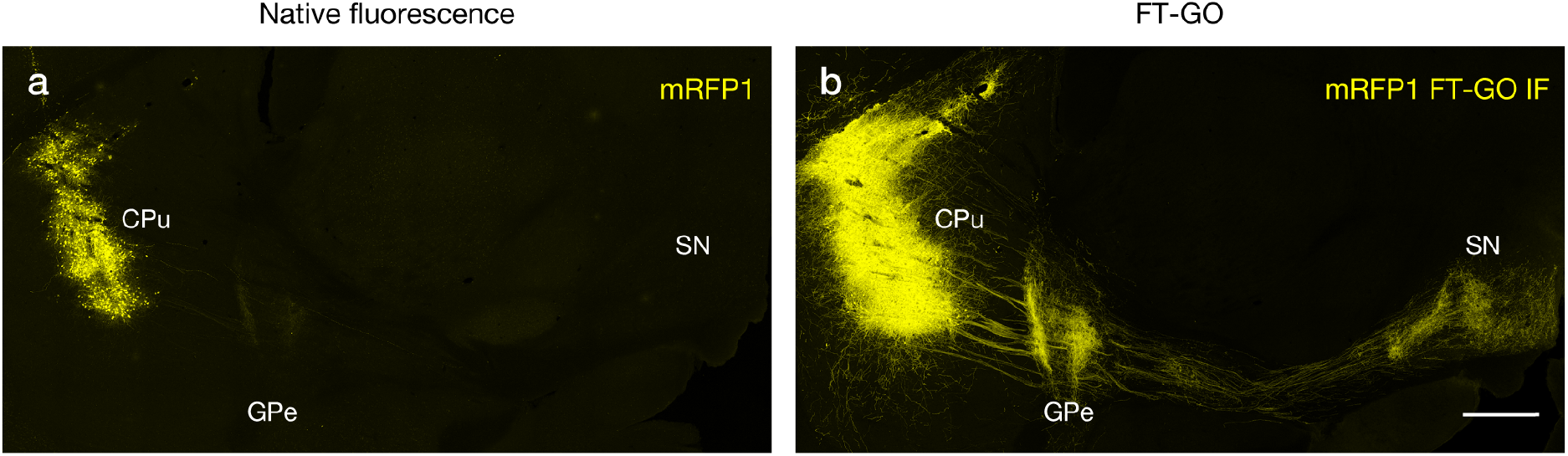
Neuroanatomical tract tracing combined an anterograde AAV tracer with FT-GO IF. **a, b**, mRFP1 native fluorescence (**a**) and RFP FT-GO IF (**b**) in mouse parasagittal brain sections with an injection of AAV2/1-SynTetOff-mRFP1 (n = 4 injection sites from 2 animals). (**a**) and (**b**) are adjacent sections. CF®568 tyramide is used for color development. CPu: caudate-putamen, GPe: external segment of the globus pallidus, SN: substantia nigra. Scale bars: 500 µm.

## Discussion

TSA is a highly sensitive enzymatic amplification method that enables detection of low-abundance targets and dramatic signal enhancement. In the present study, we reported FT-GO signal amplification as a multiplex fluorescent TSA system. Unlike conventional TSA systems, FT-GO involves POD-catalyzed deposition of FT with H_2_O_2_ produced during oxidation of glucose by glucose oxidase. We adapted FT-GO to several histochemical techniques, including IF, mRNA FISH histochemistry and neuroanatomical tract tracing with an AAV tracer. FT-GO gave strong labeling with a high signal to noise ratio in each application.

TSA has provided a remarkable increase in sensitivity of histochemical analysis, such as IHC, IF and DNA and RNA ISH histochemistry^4-7,12^. As with other TSA methods, FT-GO gave strong labeling in Rbfox3 IF (Fig. 2), and was implemented to mRNA FISH histochemistry (Fig. 6), where the increased sensitivity afforded by TSA is critical for detection of low-abundance mRNA targets^6,7^. The signal amplification efficiency of FT-GO, the enhancement of 9.7 to 34.1 fold over an indirect IF detection (Fig. 2j), is comparable to that of previous investigations with the range of 4-to 18-fold^40,41^. We utilized the high sensitivity of FT-GO to visualize fine-grain anatomy of CA and 5-HT projection systems in the mouse and marmoset brain (Fig. 3, 4, Supplementary Fig. S1) and increase detectability of an anterograde AAV tracer (Fig. 8).

The compatibility of FT-GO signal amplification to marmoset brain sections demonstrated its applicability and scalability to large-scale tissues (Fig. 4). FT-GO facilitates a high-throughput and cost-effective imaging of large-scale tissues in following ways. First, FT-GO saves image acquisition time, as strong fluorescence signal obtained by FT-GO supports reduced exposures to fluorescent and/or laser light. An improvement in image acquisition time can deliver a significant enhancement of throughput of imaging. Second, FT-GO allows to reduce the amount of primary Abs used for staining (Fig. 2, Supplementary Fig. S1). The fluorescence enhancement by FT-GO allows for using more diluted concentrations of primary Abs in comparison with conventional IF detections. This is critical in staining of large-scale tissues where a large amount of Abs is required. Additionally, inactivation of the Ab-conjugated POD with NaN_3_ enables application of multiple primary Abs in one step, avoiding need for multiple time-consuming cycles of staining with primary Abs. This is in contrast to a TSA-based multiplex IF with heat-mediated stripping of the primary/secondary Ab-POD complex that needs sequential primary Ab binding^24,25^.

Resolution is one of the main considerations for *in situ* signal amplification techniques. Although FT-GO IF visualized CA and 5-HT axons at single fiber resolution in brain regions where they diffusely innervated (Fig. 3, Supplementary Fig. S1), we did not rigorously assess the resolution of *in situ* signal amplification with FT-GO in this study. TSA can lead to blurring of signals and decreased resolution because of free diffusion of tyramide radicals before their covalent deposition on electron-rich moieties at or near the POD^42,43^. Addition of viscosity-increasing agents and/or an agent to reduce tyramide radicals lifetime to TSA reaction mixtures improves localization of fluorochromized and haptenized tyramide^42,43^. Further studies are needed to determine the resolution of *in situ* signal amplification with FT-GO.

Multiplex labeling enables detection of multiple targets on the same tissue section, allowing for examination of spatial arrangement of target molecules. For multiplexing with FT-GO, we inactivated Ab-conjugated POD by incubation with high concentration of NaN_3_ (Fig. 6, 7). Quenching Ab-conjugated POD with NaN_3_ would be effective for multiplex labeling with other TSA systems. Incubation with NaN_3_ has less impact on antigenicity, antigen-antibody binding, hybridized strands and tissue integrity, compared with other POD blocking agents such as H_2_O_2_ and low pH buffer^44^. We took advantage of the less deleterious effects of NaN_3_, and succeed in quadruple labeling combined with mRNA FISH histochemistry and IF (Fig. 7). A TSA-based multiplex IF utilizes heat-mediated stripping of the primary/secondary Ab-POD complex, which is accomplished through covalent deposition of FT on tissues, allowing for use of multiple Abs raised in the same species^24,25^. However, heat treatment might lead to degradation or enhancement of epitopes and tissue distortion. Importantly, the heat-mediated stripping, which can result in dissociation of hybridized probes, is not compatible with multiplex detection of hybridization signals.

AAV vectors carrying FP reporter genes enable robust fluorescent labeling of infected neurons with equivalent sensitivity to classical tracers, and are used as sensitive anterograde tracers for neuroanatomical mapping^35,36^. FT-GO IF labeled neuronal processes that were not detectable when observing native fluorescence, indicating that FT-GO IF increases detectability of an anterograde AAV tracer (Fig. 7). High reporter gene expression, which can be implemented by usage of strong promoters, insertion of posttranscriptional regulatory elements, preparations of high-titered virus vectors and/or longer survival times, achieves enhancement of viral-mediated fluorescence signals, increasing detectability of AAV tracers^39,45-48^. However, AAV-mediated high reporter gene expression and/or multiplicity of infection of AAV vectors could result in toxicity to neuronal, glial and neural progenitor cells^49-53^. Amplifying low tracer signals with FT-GO IF would provide a highly sensitive neuroanatomical tract tracing technique with minimal toxicity.

In summary, we have developed FT-GO as a multiplex TSA system. In the TSA system, oxidation of glucose by glucose oxidase is implemented to supply H_2_O_2_ for effective deposition of FT. FT-GO enables detection of low abundance targets as well as enhancement of fluorescence signals. Quenching of Ab-conjugated POD with high concentration of NaN_3_ permits *in situ* fluorescence detection of multiple target molecules with FT-GO. Given its simplicity, high sensitivity and high specificity, FT-GO would provide a versatile platform for histochemical analysis.

## Methods

### Animals

All animal experiments were approved by the Institutional Animal Care and Use Committees of Juntendo University (Approval No. 2021245 and 2021246) and Kyoto University (Approval No. Med Kyo 20031), and conducted in accordance with Fundamental Guidelines for Proper Conduct of Animal Experiments by the Science Council of Japan (2006). All animal experiments were performed in compliance with ARRIVE (Animal Research: Reporting In Vivo Experiments) guidelines.

Eight-to twelve-week-old male C57BL/6J mice (Nihon SLC) were used. All mice were maintained in specific pathogen-free conditions under a 12/12 hr light/dark cycle (light: 08:00–20:00) with ad libitum access to food and water.

31-month-old male and 33-months-old female common marmosets (body weight, 360 and 290 g; bred in our laboratory) were housed on a 14/10 hr light/dark cycle (light: 07:00–21:00). The animals were fed twice a day with solid food. Water was provided ad libitum. Each cage was furnished with a wooden perch, a food tray and an automatic water dispenser. The marmosets were received stereotaxic injections AAV vectors encoding GFP or RFP: analyses of these data are not included in the current study. We found no obvious abnormalities in gross brain anatomy associated with the AAV injection.

### Tissue preparation

Mice were anesthetized with an intraperitoneal injection of sodium pentobarbital (200 mg/kg; Somnopentyl, Kyoritsu Seiyaku) and perfused transcardially with 20 mL of PBS at 4ºC, followed by the same volume of 4% paraformaldehyde (PFA) (1.04005.1000, Merck Millipore) in 0.1 M phosphate buffer (PB; pH 7.4) at 4ºC. The brains of mice were removed and postfixed in the same fixative overnight (for IF) or 3 days (for FISH) at 4ºC. Perfusion fixation of marmoset brains was performed as described previously^54^.

The fixed brains were cryoprotected in 30% sucrose in 0.1M PB (for IF) or 30% sucrose in diethylpyrocarbonate (DEPC)-treated 0.1M PB at 4°C (for FISH), and cut coronally or sagittally into 30-or 40-µm-thick sections on a freezing microtome (2000R; Leica Biosystems)^13^. The sections were stored in PBS containing 0.2% NaN_3_ at 4°C (for IF) or an antifreeze solution (for FISH) that contained 30% glycerol and 30% ethylene glycol in DEPC-treated 40 mM PBS at −20°C until use.

### FT-GO reaction

Following preincubation for 10 min with an FT-GO reaction mixture that contained 10 µM FT, 3 µg/mL glucose oxidase (16831-14, Nacalai Tesque) and 1 or 2 % bovine serum albumin in 0.1 M PB, FT-GO reaction was initiated by adding ß-D-glucose (16804-32, Nacalai Tesque) into the reaction mixture at 20 µg/mL and proceeded for 30 min. The FT used were: CF®488A tyramide (92171, Biotium), CF®568 tyramide (92173, Biotium) and CF®647 tyramide (96022, Biotium).

### IF in tissue sections

Free-floating immunostaining was carried out as follows. Tissue sections were washed twice for 10 min in 0.3% Triton X-100 in PBS (PBS-X) and reacted for overnight to 3 days with primary Abs in PBS-X containing 0.12% λ-carrageenan (035-09693; Sigma-Aldrich) and 1% normal donkey serum (S30-100ML, Merck Millipore) (PBS-XCD). After washing twice for 10 min in PBS-X, the sections were then reacted for 2−4 hr with secondary Abs in PBS-XCD. The sections were washed twice for 10 min in PBS-X. Some of the sections were counterstained with NeuroTrace™ 435/455 Blue or NeuroTrace™ 530/615 Red Fluorescent Nissl Stain (1:100, N21479 or N21482, Thermo Fisher Scientific) in PBS-X. The sections were mounted onto glass slides (Superfrost micro slide glass APS-coated, Matsunami Glass) and coverslipped with 50% glycerol, 2.5% 1,4-diazabicyclo[2.2.2]octane, and 0.02% NaN_3_ in PBS (pH 7.4). Endogenous peroxidase activity was quenched with 1% H_2_O_2_ in PBS for 30 min prior to immunostaining. For multiplex FT-GO IFs, the Ab-POD conjugate was inactivated with 2% NaN_3_ in PBS for 4 hr after each round of FT deposition. All incubations were performed at 20–25ºC. Primary Abs (pAbs) and secondary Abs (sAbs) used are listed in Supplementary Table S1 and S2 online.

### Multiplex fluorescent in situ hybridization histochemistry

Multiplex FISH on free-floating sections was carried out as reported previously^16,29^, with some modifications. Briefly, tissue sections were quenched with 2% H_2_O_2_ in 0.1 M PB, acetylated with 0.25% acetic anhydride in 0.1 M triethanolamine, prehybridized in a hybridization buffer and hybridized with a mixture of 1.0 µg/ml digoxigenin (DIG)- and 1.0 µg/ml fluorescein isothiocyanate (FITC)-labeled cRNA probes. After high-stringency washes and RNase A digestion, the sections were reacted with POD-conjugated sheep (Sh) polyclonal anti-FITC Ab, Fab fragments (1:1,000, 11 426 346 910, Roche Diagnostics, RRID:AB_840257). Subsequently, the sections were washed twice for 10 min in 0.1 M Tris-HCl (pH 7.5) buffered 0.9% saline (TS7.5) containing 0.1% (v/v) Tween-20 (TNT), incubated for 10 min in 0.1 M PB and developed with first FT-GO reaction. After washes for 10 min each in 0.1 M PB and PBS, the Ab-POD conjugate was inactivated with 2% NaN_3_ in PBS for 4 hr. Following washes for 10 min each in PBS, TNT and TS7.5, the sections were then reacted with POD-conjugated Sh polyclonal anti-DIG Ab, Fab fragments (1:1,000, 11 207 733 910, Roche Diagnostics, RRID:AB_514500), washed with TNT, incubated in 0.1 M PB and developed with the second FT-GO reaction. Antisense and sense single-strand riboprobes were transcribed *in vitro* using DIG or FITC RNA labeling kit (Roche Diagnostics). cRNA probes for *Gad1*^55^ and *Slc17a7*^56^ were used.

For indirect IF for neurochemical markers, the sections processed for multiplex FISH were washed twice for 10 min in 0.1 M PB, incubated for 10 min each in PBS and PBS-X, reacted with pAbs and sAbs, and mounted onto slides as described above.

### AAV vector construction, production and injection

For the construction of pAAV2-SynTetOff-mRFP1-BGHpA, the coding sequence of mRFP1^57^ was amplified and replaced with the GFP sequence of an entry vector, pENTR1A-SV40LpA-tTAad-SYN-insulator-TRE-GFP-BGHpA^39^, through the BamHI/MluI sites. The resultant entry vector, pENTR1A-SV40LpA-tTAad-SYN-insulator-TRE-mRFP1-BGHpA, was reacted with pAAV2-DEST(f)^39^ by homologous recombination with LR clonase II (11791020, Thermo Fisher Scientific), yielding pAAV2-SynTetOff-mRFP1-BGHpA.

The AAV vector was produced and purified as described previously^58^. In brief, pAAV2-SynTetOff-mRFFP1-BGHpA and helper plasmids were transfected into HEK293T cells (RCB2202, RIKEN BioResource Research Center) with polyethylenimine (23966, Polysciences). Virus particles collected from the cell lysate and supernatant were purified by ultracentrifugation with OptiPrep (1114542, Axis-Shield) and concentrated by ultrafiltration with Amicon Ultra-15 Centrifugal Filter Unit (UFC903024, Merck Millipore). The physical titer of the virus vector (genome copies (gc)/mL) was determined by quantitative PCR with the purified virus solutions.

Stereotaxic injection of the AAV vector was carried out as reported previously^13,59^. After anesthetization with intraperitoneal injection of a mixture of medetomidine (0.3 mg/kg; Domitor®, Zenoaq), midazolam (4 mg/kg; Dormicum®, Astellas Pharma), and butorphanol (5 mg/kg; Vetorphale®, Meiji Seika Pharma), mice were placed on a stereotaxic apparatus (SR50, Narishige). Subsequently, a 0.2 µL solution of AAV2/1-SynTetOff-mRFFP1-BGHpA (1.0 × 10^12^ gc/mL) was delivered to the CPu (AP +0.5 mm, ML ±2.0 mm relative to bregma, DV –2.5 mm from the brain surface) through a glass micropipette attached to Picospritzer III (Parker Hannifin). The mice were recovered from anesthesia with intraperitoneal injection of atipamezole (1.5 mg/kg; Antisedan, Zenoaq) and maintained for seven to ten days.

### Image acquisition and processing

Images of tissue sections were acquired with a confocal laser scanning microscope (TCS SP8; Leica Microsystem) using a 5×air (HCX PL Fluotar 5x/0.15, numerical aperture [NA] = 0.15, Leica Microsystems), 10× air (HCX PL APO 10x/0.40 CS, NA = 0.40, Leica Microsystems), 16× multi-immersion (HC FLUOTAR 16x/0.60 IMM CORR VISIR, NA = 0.60, Leica Microsystems) and 25× water-immersion (HC FLUOTAR L 25x/0.95 W VISIR, NA = 0.95, Leica Microsystems) objective lens. The confocal pinhole was set to 1.0 to 5.0 Airy unit. Confocal *z*-stack images were collected at 1.0 to 30 μm intervals at 512×512 or 1024×1024 pixel resolution. Acquired images were tiled, stitched and processed to create maximum intensity projection using Leica Application Suite X software (LAS X, ver. 3.5.5.19976, Leica Microsystems) and ImageJ software^60^ (ver. 1.53, National Institutes of Health). The global brightness and contrast of the images were adjusted with ImageJ software.

### Quantitative analysis

To assess the signal amplification resulting from FT-GO reaction, the fluorescence intensity (arbitrary units [AU]) of Rbfox3 IF in layer 2/3 of the S1 was measured using ImageJ software. Rbfox3-positive cells with visible nucleoli located at approximately 3.0 µm depth from the surface were subjected to analysis. Nucleoli were identified with NeuroTrace™ 530/615 Red Fluorescent Nissl Stain.

Statistical analyses were conducted with the aid of GraphPad Prism 9 software (ver. 9.3.1 (350), GraphPad Software). Kruskal-Wallis test followed by Dunn’s *post hoc* test was used for comparisons among independent groups. Data are represented as means ± standard deviations (SDs). All tests were two-sided. The exact values of n are indicated in the corresponding figure legends. Statistical significance was set at *P* < 0.05.

## Data availability

The datasets generated during and/or analyzed during the current study and all biological materials reported in this article are available from the corresponding author on reasonable request.

## Acknowledgments

The authors thank Ayaka Ansai and Naoko Imai (Juntendo University) for their technical assistance. This study was supported by JSPS KAKENHI (JP20K07231 to K.Y.; JP21H03529 to T.F.; JP20K07743 to M.K.; JP21H02589 and JP20H05628 to M.W.; JP21H02592 to H.H.) and Scientific Research on Innovative Area “Resonance Bio” (JP18H04743 to H.H.). This study was also supported by the Japan Agency for Medical Research and Development (AMED) (JP21dm0207112 to T.F. and H.H.; JP19dm0207093 and JP18dm0207020 to T.I.), Moonshot R&D from the Japan Science and Technology Agency (JST) (JPMJMS2024 to H.H.), Fusion Oriented Research for disruptive Science and Technology (FOREST) from JST (JPMJFR204D to H.H.), Grants-in-Aid from the Research Institute for Diseases of Old Age at the Juntendo University School of Medicine (X2016 to K.Y.; X2001 to H.H.), and the Private School Branding Project.

## Author contributions

Conceptualization: K.Y., and H.H.; methodology: K.Y., and H.H.; investigation: K.Y., S.O., Y.I., K.K., K.H., T.F., M.T., K.I., M.W., and H.H.; formal analysis: K.Y., K.H., and H.H.; writing – original draft: K.Y., and H.H.; writing – review & editing: K.Y., T.F., M.K., M.W., T.I., and H.H.; project administration: H.H.; funding acquisition: K.Y., T.F., M.K., M.W., T.I., and H.H.

## Competing interests

The authors declare no competing interests.

## Notes

### Competing Interest Statement

The authors have declared no competing interest.

