## Supplementary Information for "FT-GO: a multiplex fluorescent tyramide signal amplification system for histochemical analysis"

**Title:**

Supplementary Figure S1. 5-HT innervation in the mouse brain visualized with FT-GO.

Supplementary Figure S2. Quenching of Ab-conjugated POD by incubation with $NaN_3$.

Supplementary Table S1. Primary antibodies used in the present study.

Supplementary Table S2. Secondary antibodies used in the present study.

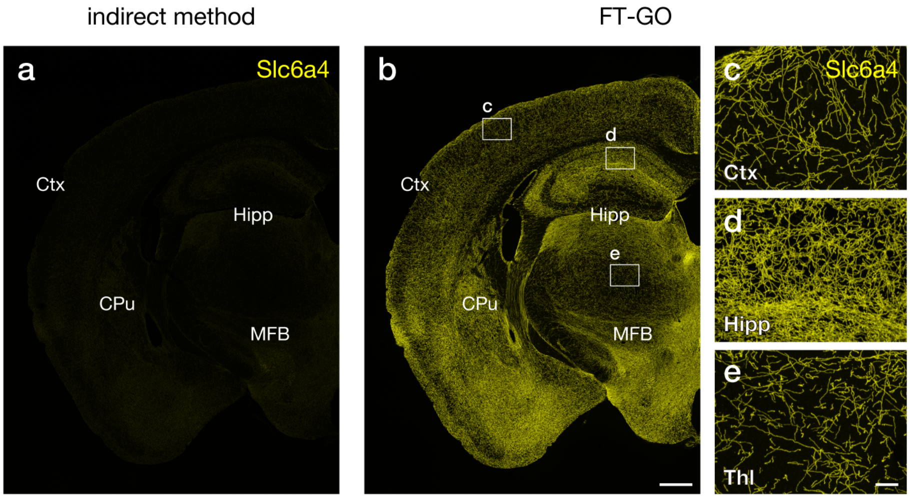


**Supplementary Fig. S1| 5-HT innervation in the mouse brain visualized with FT-GO.**
**a, b,** Slc6a4 IF in mouse brain sections visualized with an indirect (**a**) and FT-GO method (**b**) (n = 3 animals for each condition). Images are acquired with the same parameters for comparisons. Five times lower concentration of the primary Ab was applied in the FT-GO IF. **c-e.** Higher magnification images in rectangles in (**b**). CF®488A tyramide is used for color development in the FT-GO method. CPu: caudate-putamen, Ctx: cerebral cortex, Hipp: hippocampus, MFB: medial forebrain bundle, Thl: thalamus. Scale bars: 500 µm in (**b**) and 50 µm in (**e**).

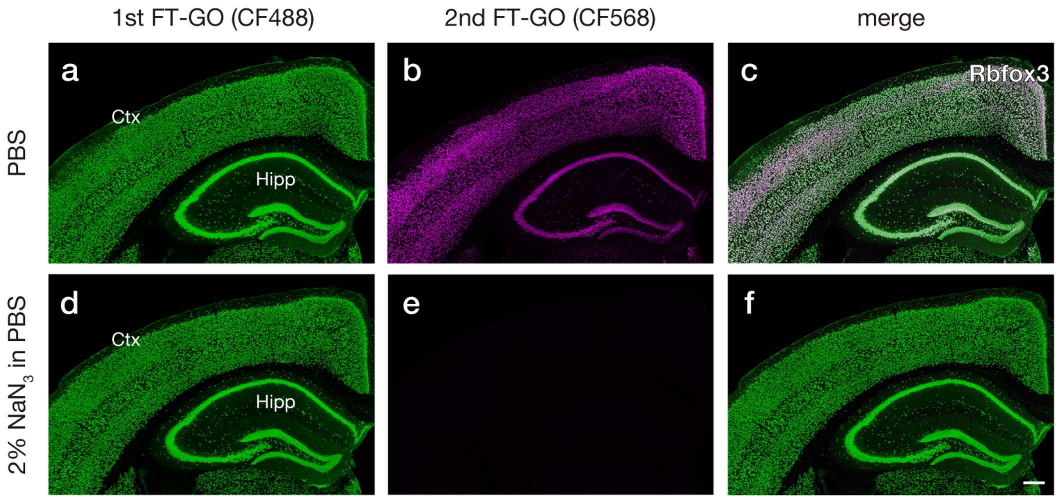


**Supplementary Fig. S2| Quenching of Ab-conjugated POD by incubation with $NaN_3$.**
**a-f,** Two rounds of FT-GO in IF for Rbfox3. Brain sections were treated with PBS (**a-c**) or 2% $NaN_3$ in PBS for 4 hr (**d-f**) between the first and second round FT-GO (n = 3 animals for each condition). First and second round of FT-GO are color-developed with CF®488A (green, **a, d**) and CF®568 tyramide (magenta, **b, e**). (**c**) and (**f**) show merged images of (**a**) and (**b**), and (**d**) and (**e**), respectively. Ctx: cerebral cortex, Hipp: hippocampus. Scale bar: 250 µm.

**Supplementary Table S1| Primary antibodies used in the present study.**

| Antigen | Host species | Source, Cat. No. | RRID | Concentration or dilution |
|---|---|---|---|---|
| Aif1 | Goat, polyclonal | FUJIFILM Wako Pure Chemical Corporation, 011-27991 | n/a | 1:5,000 |
| Gfap | Rabbit, polyclonal | Sigma-Aldrich, G9269 | AB_477035 | 1:20,000 |
| mRFP1 | Rabbit, polyclonal | Hioki et al., *J Comp Neurol*; 518, 668-686 | n/a | 0.1 µg/ml |
| proCck | Rabbit, polyclonal | Frontier Institute, CCK-pro-Rb-Af350 | AB_2571674 | 1:500 |
| Pvalb | Mouse, monoclonal | Sigma-Aldrich, P3088 | AB_477329 | 1:5,000 |
| Rbfox3 | Mouse, monoclonal | Merck Millipore, MAB377 | AB_2298772 | 1:100, 1,000, 10,000 or 100,000 |
| Rbfox3 | Mouse, monoclonal | Merck Millipore, MAB377X | AB_2149209 | 1:100, 1,000, 10,000 or 100,000 |
| Slc6a4 | Rabbit, polyclonal | Frontier Institute, HTT-Rb-Af560 | AB_2571775 | 1:1,000 (indirect detection) 1:5,000 (FT-GO detection) |
| TH | Rabbit, polyclonal | PelFreez, P40101-150 | AB_2617184 | 1:1,000 |

**Supplementary Table S2| Secondary antibodies used in the present study.**

| Antibody | Source, Cat. No. | RRID | Concentration or dilution |
|---|---|---|---|
| CF®405M Goat anti-Mouse IgG | Biotium, 20182 | AB_10557262 | 10 µg/ml |
| CF®488A Donkey anti-Mouse IgG | Biotium, 20014 | AB_10561327 | 10 µg/ml |
| CF®488A Donkey anti-Rabbit IgG | Biotium, 20015 | AB_1055966 | 10 µg/ml |
| CF®647 Donkey anti-Rabbit IgG | Biotium, 20047 | AB_10559808 | 10 µg/ml |
| POD F(ab')2 fragment Donkey anti-Goat IgG | Jackson Immuno Research, 705-036-147 | AB_2340392 | 1:200 or 500 |
| POD F(ab')2 fragment Donkey anti-Mouse IgG | Jackson Immuno Research, 715-036-151 | AB_2340774 | 1:200 or 500 |
| POD F(ab')2 fragment Donkey anti-Rabbit IgG | Jackson Immuno Research, 711-036-152 | AB_2340590 | 1:200 or 500 |
